# The role of the atypical chemokine receptor CCRL2 in myelodysplastic syndrome and secondary acute myeloid leukemia

**DOI:** 10.1101/2021.07.29.454394

**Authors:** Theodoros Karantanos, Patric Teodorescu, Brandy Perkins, Ilias Christodoulou, Christopher Esteb, Ravi Varadhan, Eric Helmenstine, Trivikram Rajkhowa, Bogdan C. Paun, Challice Bonifant, W. Brian Dalton, Lukasz P. Gondek, Alison R. Moliterno, Mark J. Levis, Gabriel Ghiaur, Richard J. Jones

**Affiliations:** Division of Hematological Malignancies, Department of Oncology, Sidney Kimmel Comprehensive Cancer Center, Johns Hopkins University Hospital, Baltimore, MD; Department of Pediatrics, Johns Hopkins University Hospital, Baltimore, MD; Division of Biostatistics and Bioinformatics, Johns Hopkins/Sidney Kimmel Comprehensive Cancer Center, Baltimore, MD; Division of Adult Hematology, Department of Medicine, Johns Hopkins University, Baltimore MD

## Abstract

The identification of new molecular pathways supporting the growth of myelodysplastic syndrome (MDS) stem and progenitor cells is needed to understand clinical variation and develop targeted therapies. Within myeloid malignancies, men have worse outcomes compared to women, suggesting male sex hormone driven effects in malignant hematopoiesis. The androgen receptor promotes the expression of five granulocyte-colony factor receptor regulated genes. Among them, *CCRL2* encodes an atypical chemokine receptor that regulates cytokine signaling in differentiated granulocytes but its role in myeloid malignancies is unknown. Our study revealed that CCRL2 is upregulated in stem and progenitor cells from patients with MDS and secondary acute leukemia. CCRL2 knockdown suppressed the growth and clonogenicity of MDS92 and MDS-L cells *in vitro* and *in vivo*. Moreover, CCRL2 knockdown significantly suppressed the phosphorylation of JAK2, STAT3, and STAT5 in MDS cells. CCRL2 co-precipitated with JAK2 and its suppression decreased the interaction of JAK2 with STAT proteins. Cell lines expressing JAK2V617F showed less effect of CCRL2 knockdown on growth and clonogenicity compared to those expressing wild type. However, the selective JAK2 inhibitor fedratinib potentiated the effects of CCRL2 knockdown in MDS and leukemia cells expressing both wild type JAK2 and JAK2V617F. In conclusion, our results implicate CCRL2 as a mediator of MDS and secondary acute leukemia cell growth, in part through JAK2/STAT signaling.

## INTRODUCTION

Myelodysplastic syndromes (MDS) arise from hematopoietic stem and progenitor cells that acquire somatic mutations (*1*). Although they can respond to chemotherapy and hypomethylating agents, MDS are only cured by allogeneic transplantation (*1, 2*). Oncogenic pathways involving granulocyte colony stimulating factor receptor (GCSF-R) and JAK2/STAT signaling have been implicated in the progression of myeloproliferative neoplasms (MPN), MDS/MPN and acute myeloid leukemia (AML) (*3, 4*), but their roles in MDS are less well understood. A deeper understanding of the molecular pathways providing survival and growth advantage to MDS cells over healthy hematopoietic cells is critical for improving the outcomes of patients with these diseases.

Recent studies support that men with myeloid neoplasms generally exhibit higher disease burdens and experience worse outcomes compared to women independent of other known prognostic markers (*5–7*). However, the underlying molecular mechanisms responsible for these observations are unclear. Hormonal receptors are expressed in hematopoietic cells (*8*) and estrogen receptor signaling induces the apoptosis of malignant stem cells (*9*). The androgen receptor (AR) promotes normal granulopoiesis by activating in myeloid precursors the GCSF-R with five of the most upregulated GCSF-R target genes being: *CCRL2*, *C3AR1*, *GYK*, *SOCS3,* and *FLOT2* (*10*). Among these, *C3AR1* is overexpressed in AML cells (*11*), and this has been associated with worse outcomes in AML patients (*12*). Similarly, *FLOT2* promotes the homing of chronic myeloid leukemia initiating cells in a mouse model (*13*), and *SOCS3* suppresses the CD33-mediated inhibition of cytokine induced proliferation of AML cells (*14*).

*CCRL2* encodes for one of the atypical chemokine receptors (*15*), and is expressed in granulocytes, monocytes, and NK cells, playing a role in the migration of these cells to sites of inflammation (*16, 17*). CCRL2 is required for the activation of CXCR2, the chemokine receptor for IL-8 (*18*). Of note, the IL-8/CXCR2 pathway is critical for the growth of MDS and secondary AML (sAML) cells (*19*). However, little is known about the regulation and signaling role of CCRL2 in myeloid neoplasms.

Here we show that CCRL2 is upregulated in stem and progenitor cells from MDS patients. It promotes the growth and clonogenicity of MDS and sAML cells, and activates JAK2/STAT signaling. CCRL2 knockdown inhibits the growth of MDS and sAML cells, and potentiates the effect of the JAK2 inhibitor fedratinib.

## RESULTS

### *CCRL2* is upregulated in MDS and MDS-related AML

Expression of the most AR upregulated GCSF-R-target genes (*10*) in healthy stem and progenitor cells, as well as in sub-types of AML and MDS, was analyzed using the BloodSpot data set (211434_s_at) (*20*). AML/MDS cells express higher levels of *C3AR1*, *CCRL2* and SOCS3 compared to healthy cells (**Figure 1A**). *CCRL2* was the only one of the five genes found to be upregulated in MDS and AML with MDS-related chromosomal changes (which will be referred to as sAML), compared to de-novo AMLs (**Figure 1A**).

**Figure 1.**
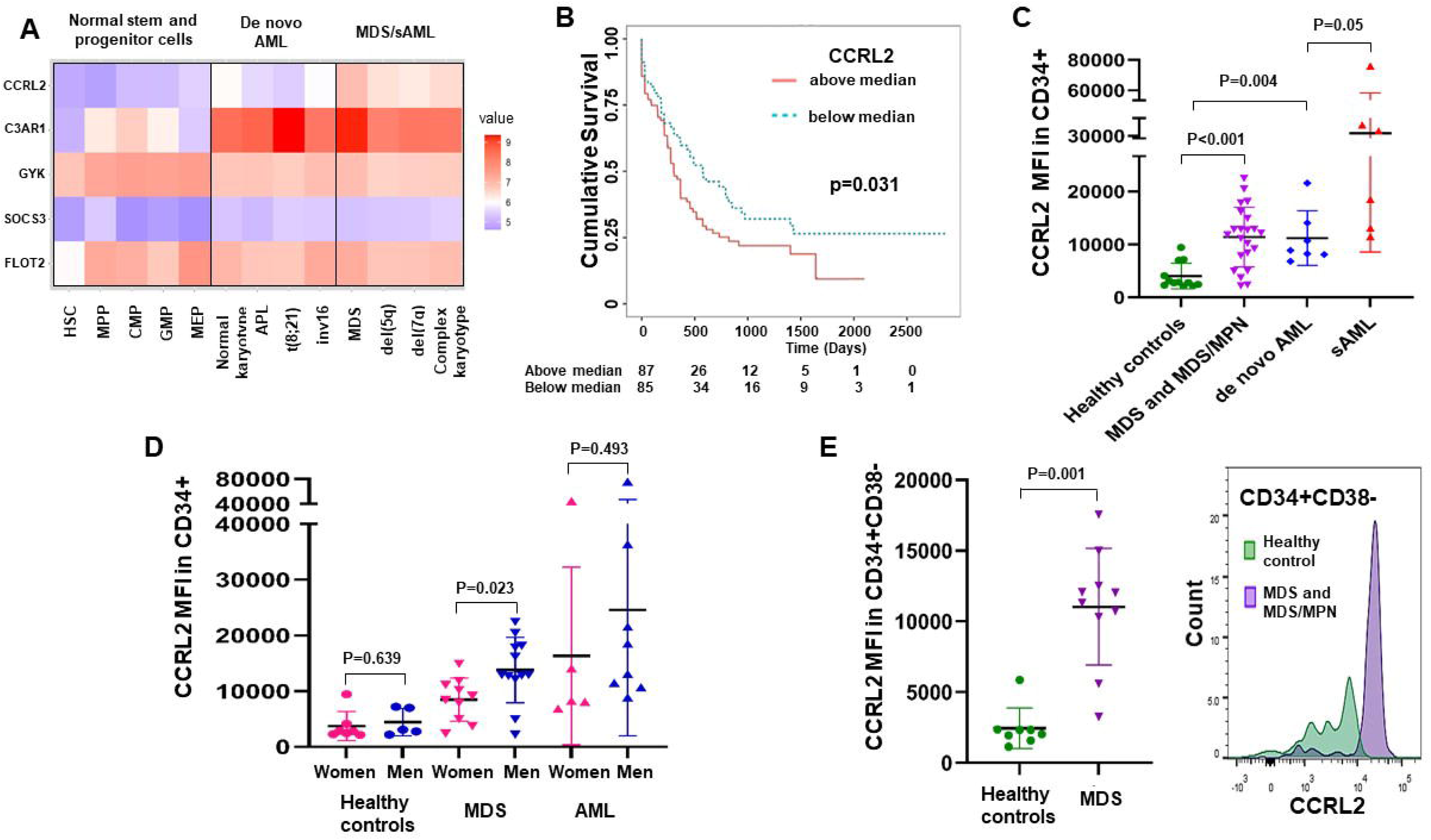
CCRL2 is upregulated in MDS and sAML cells. **A.** Comparison of the mRNA levels of the five AR regulated genes *CCRL2*, *C3AR1*, *GYK*, *SOCS3* and *FLOT2* between healthy stem and progenitor cells, de novo AML (AMLs without MDS-related chromosomal abnormalities), MDS, and sAML (AMLs with MDS-related chromosomal abnormalities) based on data extracted from the blood.spot database. *CCRL2* is the only gene that is upregulated in MDS and sAML compared to both healthy cells and de novo AMLs (*P*<0.001). **B**. AML patients with high CCRL2 expression (based on median expression) have significantly worse overall survival (P=0.031) compared to AML patients with low CCRL2 expression based on the TCGA dataset. **C.** Analysis of the CCRL2 protein levels by flow cytometry showed that CD34+ cells from MDS and MDS/MPN patients express higher levels of CCRL2 (*P*<0.001) compared to CD34+ cells from healthy controls. CD34+ blasts from patients with sAML express higher levels of CCRL2 compared to CD34+ blasts from patients with de novo AML (*P*=0.05). **D.** Healthy controls and MDS/AML patients were stratified by sex demonstrating that men with MDS and MDS/MPN express higher levels of CCRL2 compared to women with these diseases (*P*=0.023). **E.** CCRL2 expression is higher in CD34+CD38-cells from MDS patients compared to CD34+CD38-cells from healthy individuals (*P*=0.001). MFI: mean fluorescent intensity. Graphs show the mean value and standard deviation of the mean value.

Analysis of AML patient overall survivals based on median gene expression measured in the Cancer Genome Atlas (TCGA) dataset (211434_s_at) (*20*) showed that *CCRL2* expression was the only identified gene among the most AR-upregulated genes with a significant survival impact. Patients whose AMLs exhibiting CCRL2 expression above the mean had poorer overall survivals (**Figure 1B**).

### CCRL2 is upregulated in progenitor cells from MDS and AML patients, especially males

To confirm the results from publicly available databases, the expression of CCRL2 in CD34+ and CD34+CD38-bone marrow cells from 12 healthy controls and 22 patients with either MDS or MDS/myeloproliferative neoplasms (MPN) as well as CD34+ blasts from 13 AML patients was analyzed by flow cytometry (**Supplementary Figure 1A, B)**. Patient and healthy controls demographic data are summarized in **Supplementary Table 1.**

CCRL2 was upregulated in CD34+ cells from MDS and MDS/MPN patients compared to healthy controls (**Figure 1C, Supplementary Figure 1C**). sAML CD34+ cells also expressed higher CCRL2 compared to de novo AML CD34+ cells (**Figure 1C**). No significant difference was identified between MDS patients with <5% or ≥ 5% blasts, or between MDS and MDS/MPN patients (**Supplementary Figure 1D**). Although CCRL2 is upregulated in CD34+ cells from the entire cohort of MDS and MDS/MPN patients, it was significantly higher in men with MDS and MDS/MPN compared to women (**Figure 1D**). Patients with myeloid malignancies were significantly older than healthy controls (P<0.001 for MDS and MDS/MPN and P=0.020 for AML), and as there was not significant age overlap between the groups, age-adjustment could not be done. However, the CCRL2 expression in CD34+ cells showed a negative correlation with age among healthy controls (Coef −128.98, P=0.018) suggesting that older individuals may have lower CCRL2 expression in their CD34+ cells. The analysis of 8 healthy controls and 10 MDS patients (**Supplementary Table 2**) revealed that CCRL2 is also upregulated in CD34+CD38-cells from MDS patients compared to healthy controls (**Figure 1E**). Among the 10 MDS patients, CCRL2 expression in CD34+CD38-was also higher in men compared to women (**Supplementary Figure 1E**).

### CCRL2 influences the growth, clonogenicity and differentiation of MDS cell lines

The expression of CCRL2 protein in AML and MDS cell lines and sorted CD34+ cells from 3 healthy individuals was assessed by flow cytometry. MDS-L cells, a leukemic subline derived from the MDS92 cell line established from a patient with MDS with 5q deletion, monosomy 7 and *NRAS* mutation (*21, 22*), expressed the highest levels of CCRL2 from a panel of 6 MDS and AML cell lines (**Figure 2A**). This finding was confirmed by western blotting (**Supplementary Figure 2A**).

**Figure 2.**
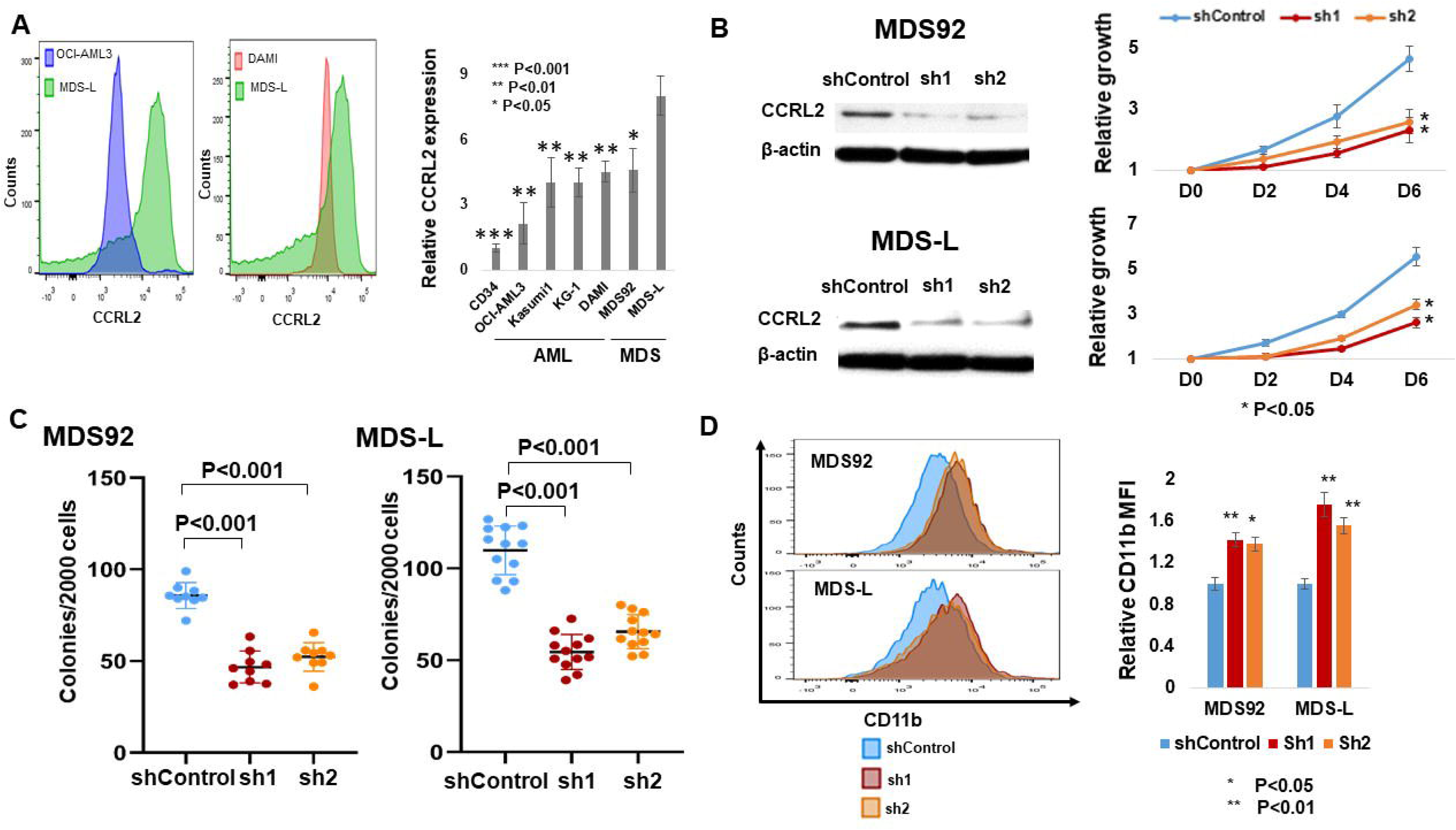
CCRL2 promotes the growth and clonogenicity of MDS cell lines. **A.** MDS-L cells express higher levels of CCRL2 compared to CD34+ cells from 3 separate healthy controls in culture (<0.001), de novo AML cell lines (OCI-AML3 (*P*=0.002) and Kasumi-1 (*P*=0.009)), sAML cell lines (KG-1 (*P*=0.004) and DAMI (*P*=0.004)) and MDS92 cells, n=3. **B**. CCRL2 was knocked down in MDS92 and MDS-L by transduction with 2 different shRNAs (sh1 and sh2) targeting CCRL2 using empty vector as control. CCRL2 knockdown suppressed the growth rate at 2 days (MDS92: *P*=0.001 with sh1 and *P*=0.034 with sh2, MDS-L: *P*=0.002 with sh1 and *P*=0.028), 4 days (MDS92: *P*=0.007 with sh1, *P*=0.028 with sh2, MDS-L: *P*<0.001 with sh1 and *P*=0.003 with sh2) and 6 days (MDS92: *P*=0.001 with sh1 and *P*=0.002 with sh2, MDS-L: *P*=0.003 with sh1 and *P*=0.010 with sh2) following puromycin selection, n=3. **C.** Evaluation of surface CD11b expression in MDS cell lines following cell culture for 6 days after puromycin selection revealed that CCRL2 knockdown increases the CD11b expression in MDS92 (*P*=0.002 with sh1 and *P*=0.009 with sh2) and MDS-L cells (*P*=0.004 with sh1 and *P*=0.003 with sh2), n=3. **D.** CCRL2 knockdown suppresses the clonogenic capacity of MDS92 (*P*<0.001 with both sh1 and sh2) and MDS-L (*P*<0.001 with both sh1 and sh2), n=9 for MDS92 and n=12 for MDS-L. Results show the mean value and standard deviation of the mean value.

To further characterize the role of CCRL2 in the survival and growth of MDS cells, MDS92 and MDS-L cells were transduced with either empty vector or two shRNA targeting CCRL2 (sh1CCRL2 and sh2CCRL2). CCRL2 knockdown was confirmed at both the RNA (**Supplementary Figure 2B**) and protein level (**Figure 2B, Supplementary Figure 2B**). MDS92 and MDS-L cells with suppressed CCRL2 exhibited significantly decreased growth rate at 2, 4 and 6 days (**Figure 2B**). Similarly, the clonogenic capacity of MDS92 and MDS-L transduced with shCCRL2 was suppressed (**Figure 2C**). To determine if induction of differentiation contributes to the diminished clonogenicity, the effect of CCRL2 knockdown in MDS cells differentiation was assessed. CD11b surface expression was used as it is frequently altered in MDS (*23*) and its levels have been previously determined in MDS-L cells (*24*). MDS cells transduced with shCCRL2 showed an increase of CD11b expression (**Figure 2D**).

### CCRL2 promotes the growth of MDS-L cells *in vivo*

The impact of CCRL2 knockdown in an MDS-L xenograft mouse model was evaluated. MDS-L shCCRL2 and control cells were cultured for 10 days in puromycin and the suppression of CCRL2 expression was assessed at the RNA and protein levels (**Figure 3A**) (**Supplementary Figure 2B**). The cells were then transduced with a GFP+/luciferase+ dual reporter retrovirus and GFP+ cells were injected intravenously in NSG-hSCF/hGM-CSF/hIL3 (NSGS) mice 48 hours after intraperitoneal injection of Clophosome-A clodronate liposomes (*25*). MDS-L shCCRL2 cells demonstrated a slower proliferative rate than control cells (**Figure 3B**). No significant differences in peripheral blood counts were observed (**Supplementary Figure 2C**). At day 78, all mice were sacrificed and a higher percentage of human CD45+ cells was found in the bone marrow (**Figure 3C**) and the spleen (**Supplementary Figure 2C**) of control mice compared to those that received MDS-L shCCRL2.

**Figure 3.**
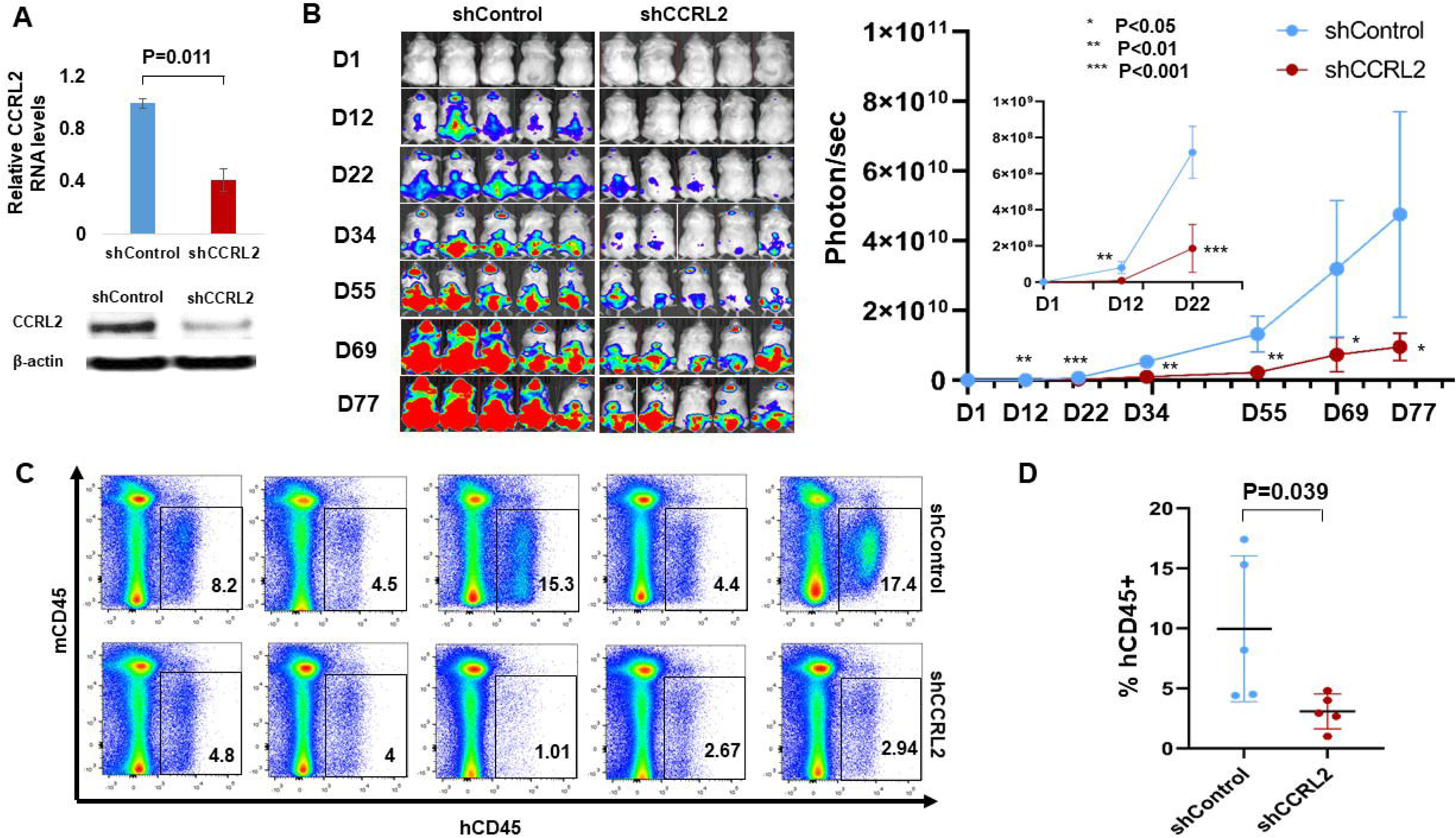
CCRL2 knockdown suppresses the engraftment and growth of MDS-L cells in NSGS mice. **A.** MDS-L cells transduced with shControl or shCCRL2 shRNAs were treated for 10 days with puromycin, and their CCRL2 expression was assessed bt RNA and protein. MDS-L cells transduced with shCCRL2 expressed significantly lower levers of CCRL2 RNA (*P*=0.011) and protein (*P*=0.006). **B.** MDS-L cells with normal or suppressed CCRL2 were transduced with a GFP+/Luciferase dual reporter retrovirus and injected to NSGS mice. The bioluminescence signal in mice injected with MDS-L with suppressed CCRL2 was significantly lower at 12 (*P*=0.0016), 22 (*P*=0.0002), 34 (*P*=0.0010), 55 (*P*=0.0017), 69 (*P*=0.0259) and 77 days (*P*=0.0212) after the injection. The bioluminescence signaling of the two group of mice is shown in a logarithmic scale. **C.** MDS-L burden measured by percentage of human CD45+ cells in the bone marrow of NSGS mice was significantly lower in mice injected with MDS-L cells with suppressed CCRL2 (*P*=0.039). Results depict the mean value and standard deviation of the mean value.

### CCRL2 affects JAK2/STAT signaling in MDS cells

CCRL2 appears to be involved in the activation of cytokine-mediated pathways such as IL-8/CXCR2, which in turn activates AKT and ERK signaling in differentiated myeloid cells (*18*). Moreover, MDS92 and MDS-L cells are dependent on IL-3, which signals through JAK2/STAT (*21, 22*), suggesting that this cytokine-mediated pathway is also critical for the growth of these cells. Thus, the effect of CCRL2 knockdown in the phosphorylation of AKT, ERK, JAK2 and STAT proteins was assessed.

The knockdown of CCRL2 in MDS cells had no impact on the phosphorylation of AKT or ERK (**Supplementary Figure 3A, B**). However, CCRL2 knockdown suppressed the phosphorylation of JAK2, STAT3 and STAT5 in both MDS92 and MDS-L cells (**Figure 4A**), (**Supplementary Figure 3B**). The effect of CCRL2 on the expression of JAK2/STAT target genes in MDS cells was also evaluated. Knockdown of CCRL2 in MDS92 and MDS-L cells decreased the expression of MYC, PIM1, BCL2, MCL1, and DNMT1 at the RNA (**Figure 4B**) and protein levels **(Supplementary Figure 3C, D),** suggesting that these genes are regulated by CCRL2 likely at the transcriptional level.

**Figure 4.**
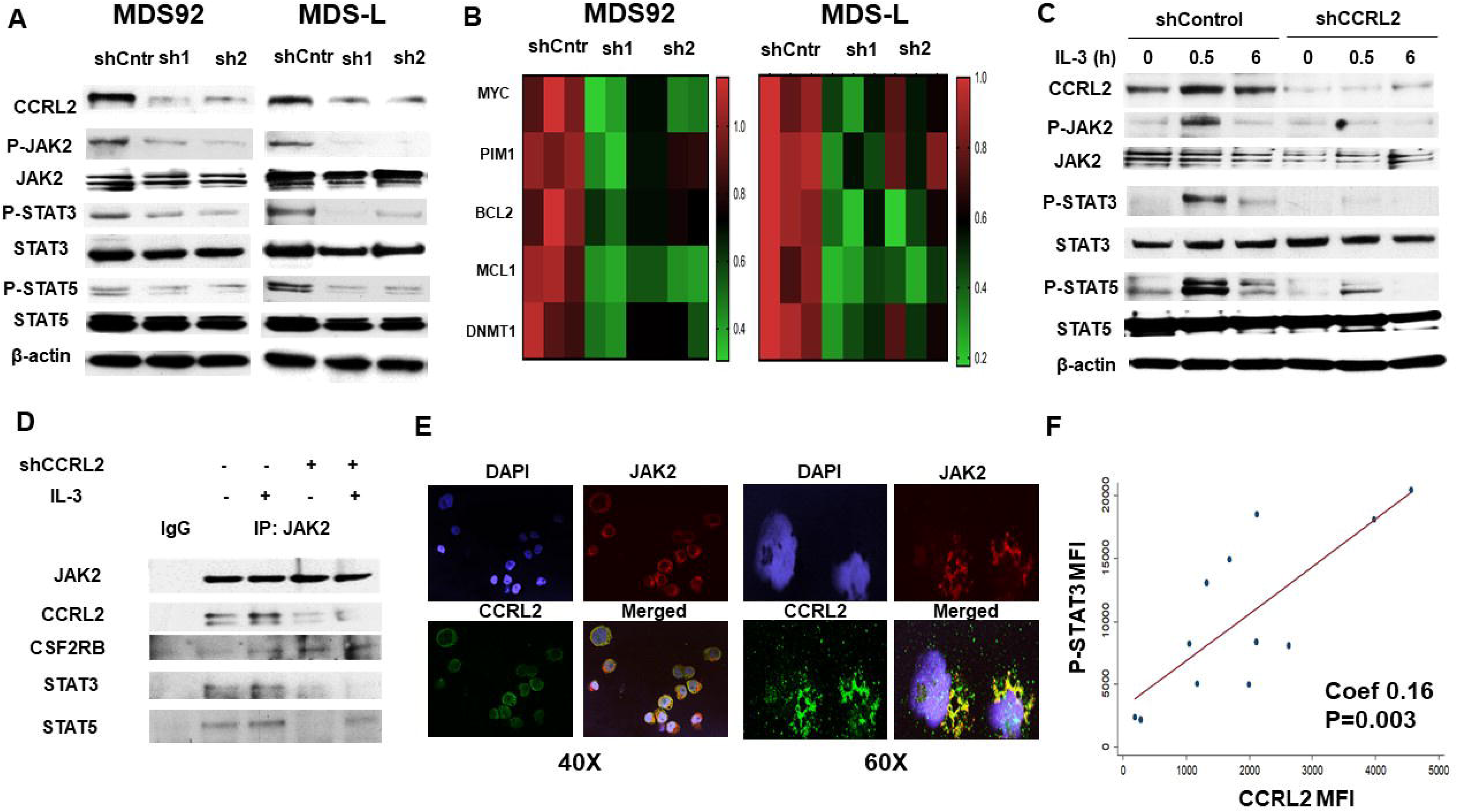
CCRL2 affects JAK2/STAT signaling in MDS cells. **A.** Representative western blotting showing the effect of CCRL2 knockdown by two different shRNAs (sh1 and sh2) on the phosphorylation of JAK2 (Tyr1007/1008), STAT3 (Tyr105) and STAT5 (Tyr694) in MDS92 and MDS-L with empty vector (shControl) as control. **B.** CCRL2 knockdown with two different shRNAs (sh1 and sh2) decreases the RNA levels of the JAK2/STAT target genes: *MYC* (MDS92: *P*=0.001 with sh1 and *P*=0.004 with sh2, MDS-L: *P*=0.002 with sh1 and *P*=0.020 with sh2), PIM1 (MDS92: *P*=0.002 with sh1 and *P*=0.015 with sh2, MDS-L: *P*=0.003 with sh1 and *P*=0.080 with sh2), BCL2 (MDS92: *P*=0.031 with sh1 and *P*=0.071 with sh2, MDS-L: *P*=0.006 with sh1 and *P*=0.010 with sh2), MCL1 (MDS92: *P*=0.036 with sh1 and *P*=0.052 with sh2, MDS-L: *P*=0.004 with sh1 and *P*=0.036 with sh2) and DNMT1 (MDS92: *P*=0.008 with sh1 and *P*=0.013 with sh2, MDS-L: *P*=0.010 with sh1 and *P*=0.047 with sh2), n=3. **C.** Western blotting showing that CCRL2 knockdown suppresses the phosphorylation of JAK2, STAT3 and STAT5 at 30 minutes and 6 hours of IL-3 20 ng/ml treatment following 48 hours of IL-3 starvation. **D.** Co-immunoprecipitation assay showing that CCRL2 precipitates with JAK2, and that CCRL2 knockdown does not affect the interaction between JAK2 and the common β signal transducing subunit of CD123 (CSF2RB) but decreases the interaction between JAK2 and STAT3/5 proteins. **E.** Representative images from immunofluorescence staining (40X and 60X magnification) showing localization of CCRL2 (green) in the cytoplasm and membrane of MDS-L cells. Confocal microscopy reveals areas of co-localization with JAK2 (red). **F.** The mean fluorescent intensity (MFI) of phosphorylated STAT3 (P-STAT3) is positively associated with the MFI of CCRL2 in CD34+ cells from MDS patients and CD34+ blasts from AML patients (Coef 0.16, *P*=0.003), n=12.

To further study JAK2/STAT regulation by CCRL2, MDS-L cells were deprived of IL-3 for 48 hours and then were treated with 20 ng/ml IL-3. CCRL2 knockdown suppressed IL-3 mediated JAK2 auto-phosphorylation, as well as the phosphorylation of STAT3 and STAT5 **(Figure 4C)**, suggesting that CCRL2 has a role in the IL-3 receptor (CD123), JAK2 and STAT interaction. Co-immunoprecipitation showed that CCRL2 is associated with JAK2, and CCRL2 knockdown suppresses the JAK2/STAT interaction following IL-3 treatment **(Figure 4D)**. In contrast, CCRL2 knockdown did not affect the interaction between JAK2 and the common β signal transducing subunit of the IL-3 receptor (CSF2RB) **(Figure 4D)**. Immunofluorescence analysis revealed the localization of CCRL2 in the membrane and cytoplasm of MDS-L cells and confocal microscopy showed areas of co-localization with JAK2 (**Figure 4E**).

CCRL2 and STAT3 and STAT5 phosphorylation was assessed by flow cytometry in primary CD34+ cells from 9 healthy controls, 8 patients with MDS, and 4 AML patients (**Supplementary Figure 3D**). CCRL2 expression was positively associated with the phosphorylation of STAT3 (**Figure 4F**), but not with the phosphorylation of STAT5 (**Supplementary Figure 3E**).

### CCRL2 has differential effects on JAK2 wild-type and mutated cell lines

Given the observation that CCRL2 impacts JAK2/STAT signaling, the effect of CCRL2 knockdown was assessed in two erythroleukemia cell lines with and without mutational JAK2 activation. GM-CSF-dependent TF-1 cells are *JAK2* wild type, while DAMI cells carry a *JAK2V617F* mutation and are cytokine independent (*26, 27*). CCRL2 knockdown significantly decreased the growth of TF-1 cells in the presence and absence of GM-CSF (**Figure 5A**) while its effect on the DAMI cells growth was less prominent (**Figure 5A**). Similarly, CCRL2 knockdown had a more dramatic effect on the clonogenicity of TF-1 cells in the presence and particularly in the absence of GM-CSF compared to DAMI cells (**Figure 5B**). Finally, the expression of CD71 was measured by flow cytometry to assess the induction of erythroid differentiation (*28*). CCRL2 knockdown also augmented the induction of erythroid differentiation in TF-1 cells compared to DAMI cells (**Figure 5C**). CCRL2 knockdown suppressed JAK2 phosphorylation in both cell lines, but STAT3 and STAT5 phosphorylation was decreased only in TF-1 (**Figure 5D**).

**Figure 5.**
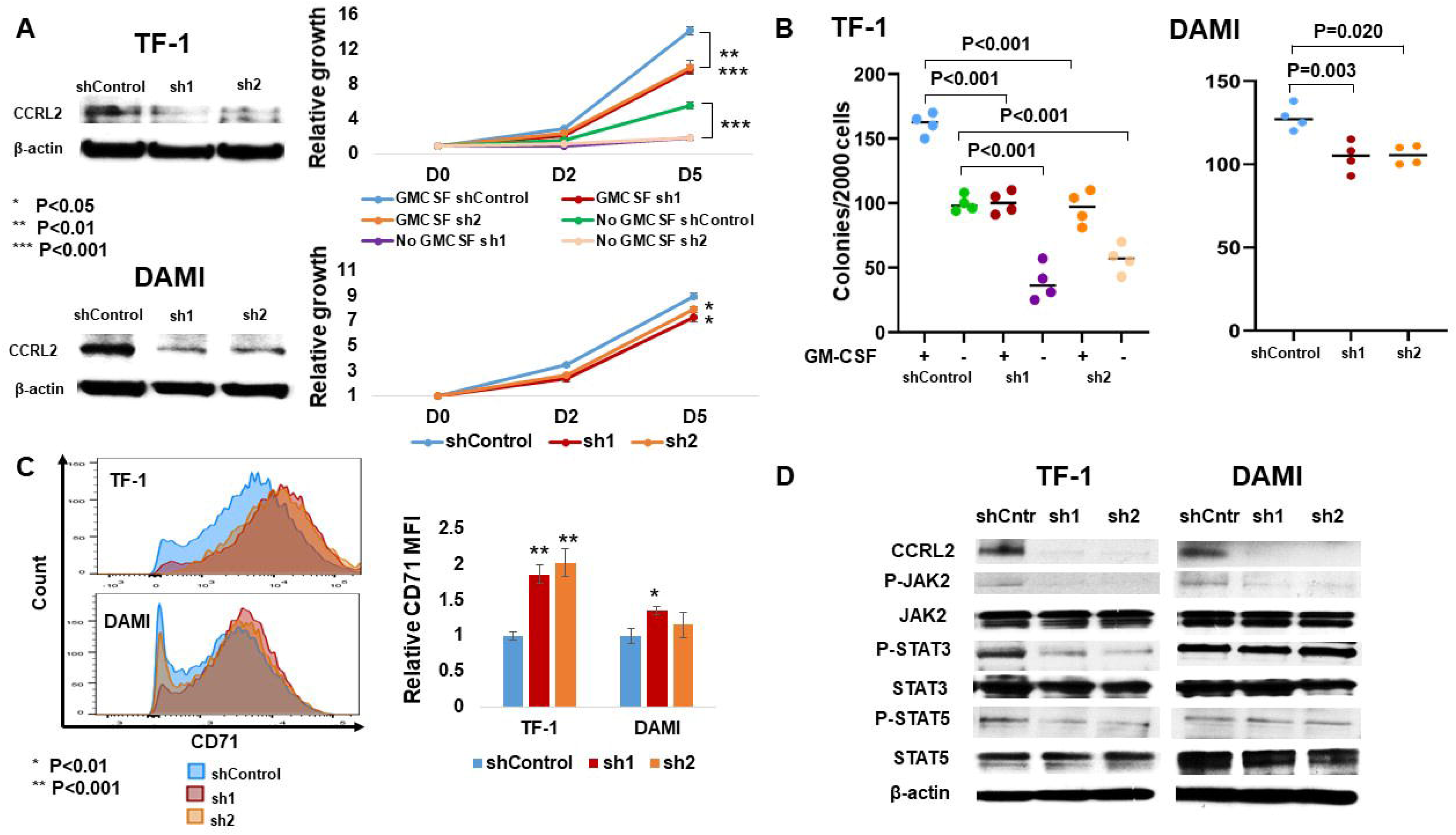
CCRL2 knockdown affects the growth and clonogenicity of erythroleukemia cell lines. **A.** CCRL2 knockdown with two different lentiviruses significantly suppresses the growth of TF-1 cells in the presence (*P*<0.001 with sh1 and *P*=0.004 with sh2) and absence of GM-CSF (*P*<0.001 with both sh1 and sh2). CCRL2 knockdown with two different lentiviruses suppresses the growth of DAMI cells (*P*=0.031 with sh1 and *P*=0.032 with sh2). **B.** CCRL2 knockdown with two different lentiviruses decreases the colony formation of TF-1 cells in the presence (*P*<0.001 with both sh1 and sh2) and absence of GM-CSF (*P*<0.001 with both sh1 and sh2). CCRL2 knockdown with two different lentiviruses decreases the colony formation of DAMI cells (*P*=0.003 with sh1 and *P*=0.020 with sh2) at a lower extent compared to TF-1 cells. **C.** CCRL2 knockdown with two different lentiviruses increases the CD71 expression in TF-1 cells (*P*<0.001 with both sh1 and sh2). CCRL2 knockdown increases the expression of CD71 at a significantly lower extent in DAMI cells (*P*=0.004 with sh1 and *P*=0.074 with sh2). **D.** Western blotting showing the effect of CCRL2 knockdown in the JAK2/STAT signaling in TF-1 and DAMI cells. CCRL2 knockdown suppresses the phosphorylation of JAK2, STAT3 and STAT5 in TF-1 cells. CCRL2 knockdown decreases the phosphorylation of JAK2 but does not affect the phosphorylation of STAT3 and STAT5 in DAMI cells.

### CCRL2 knockdown potentiates the effect of JAK2 inhibition

JAK2 inhibition induces the auto-phosphorylation of JAK2, which has been associated with a withdraw phenomenon caused by activation of downstream signaling in myelofibrosis patients (*29–31*) suggesting that inhibition of JAK2 auto-phosphorylation could increase the activity of JAK2 inhibition. The effect of CCRL2 knockdown on the efficacy of JAK2 inhibition by the selective JAK2 inhibitor fedratinib (*32*) was studied. MDS/AML cell lines were treated with fedratinib and the methylene tetrazolium (MTT) assay was used to assess the half maximal inhibitory concentration dose (IC50) of the drug at 72 hours of treatment. MDS92 and MDS-L had the lowest IC50 compared to all the other lines, including JAK2-mutated DAMI cells and erythroleukemia TF-1 cells (**Supplementary Figure 4A, B**).

MDS-L, TF-1, and DAMI cells with normal or suppressed CCRL2 expression were treated with PBS or 0.25 μΜ fedratinib, a dose that is lower than the IC50 dose for these 3 lines. CCRL2 knockdown further suppressed the growth of cells treated with 0.25 μΜ fedratinib at 4 days of treatment (**Figure 6A-C**). Similarly, the anti-clonogenic effect of fedratinib in MDS-L, TF-1 and DAMI cells was further increased by CCRL2 knockdown (**Figure 6D-F**).

**Figure 6.**
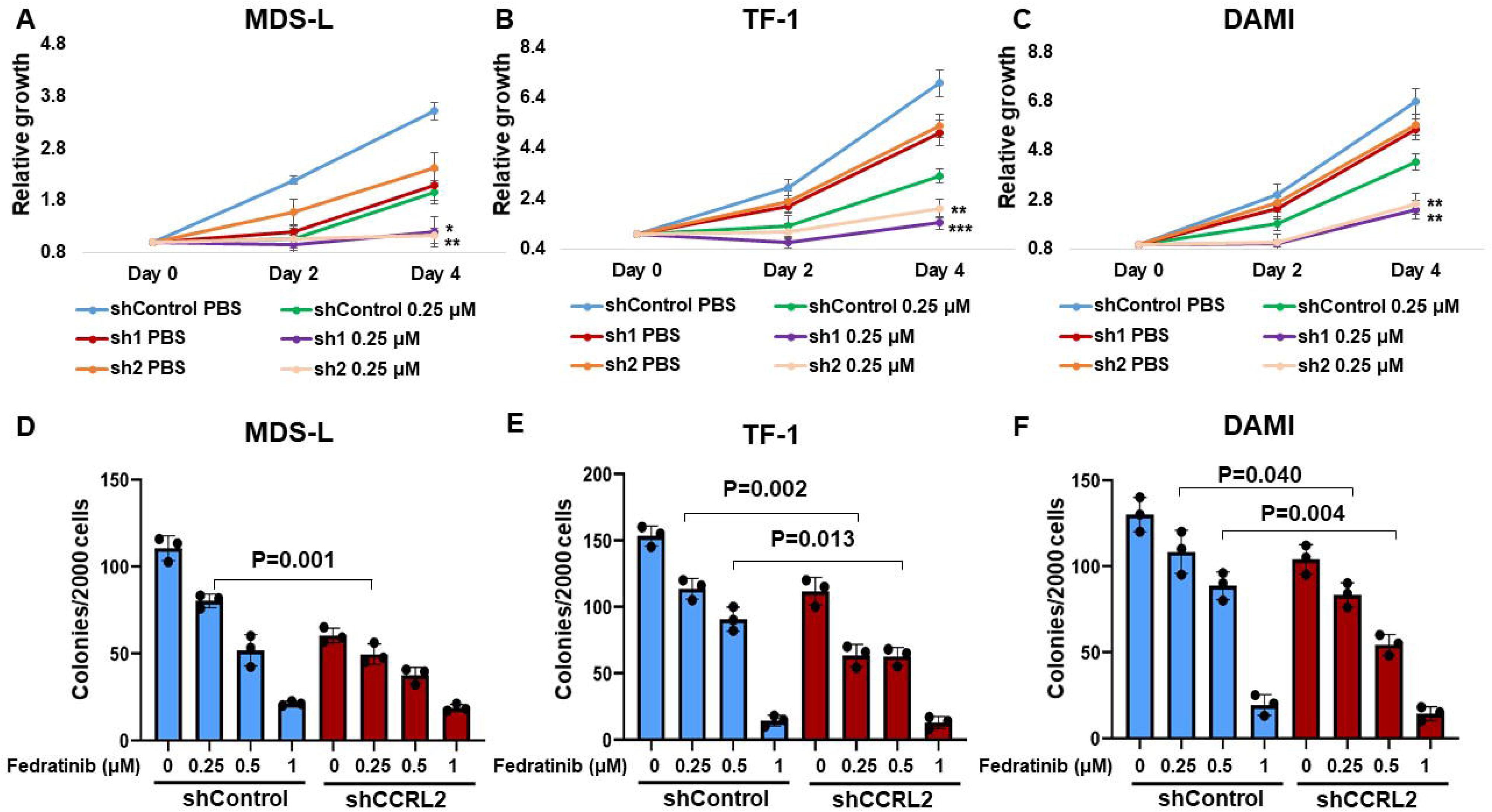
CCRL2 knockdown increases the efficacy of JAK2 inhibition by the selective inhibitor fedratinib. **A.** CCRL2 knockdown increases the anti-proliferative effect of 0.25 μΜ fedratinib (*P*=0.023 with sh1 and *P*=0.005 with sh2) in MDS-L cells. **B.** CCRL2 knockdown increases the anti-proliferative effect of 0.25 μΜ fedratinib in TF-1 cells (*P*<0.001 with sh1 and *P*=0.007 with sh2). **C.** CCRL2 knockdown increases the anti-proliferative effect of 0.25 μΜ fedratinib (*P*=0.003 with sh1 and *P*=0.005 with sh2) in DAMI cells. **D.** CCRL2 knockdown increases the anti-clonogenic effect of 0.25 μΜ fedratinib (*P*=0.001) in MDS-L cells. **E.** CCRL2 knockdown increases the anti-clonogenic effect of 0.25 μΜ (*P*=0.002) and 0.5 μΜ fedratinib (*P*=0.013) in TF-1 cells. **F.** CCRL2 knockdown increases the anti-clonogenic effect of 0.25 μΜ (*P*=0.004) and 0.5 μΜ fedratinib (*P*=0.040) in DAMI cells. * *P*<0.05, ** *P*<0.01, *** *P*<0.001.

## DISCUSSION

Our findings suggest that one of the most AR-upregulated genes implicated in the later stages of normal granulopoiesis, CCRL2, is overexpressed in stem and progenitor cells from MDS patients and CD34+ AML blasts. Knockdown of CCRL2 inhibited the growth and clonogenicity of MDS cell lines both *in vitro* and *in vivo*. Downregulation of CCRL2 decreased the phosphorylation of JAK2/STAT and the expression of STAT target genes. CCRL2 modulated IL-3-mediated phosphorylation of JAK2/STAT and the physical interaction between JAK2 and STAT3/5 proteins. CCRL2 knockdown inhibited the growth of the erythroleukemia cell lines TF-1 and DAMI, with effect on DAMI cells likely mitigated by constitutively active JAK2/STAT signaling due to the JAK2V617F mutation. Further, CCRL2 knockdown potentiated the anti-proliferative and anti-clonogenic effect of the selective JAK2 inhibitor fedratinib in MDS-L, TF-1 and DAMI cells.

Men with chronic myeloid neoplasms have worse outcomes compared to women (*5–7*). Yet, the underlying biology, including the role of AR signaling, remains poorly understood (*33*). Chuang et al reported that mice lacking AR develop severe neutropenia with deficient granulopoiesis, characterized by downregulation of five GCSF-R target genes (*10*). Based on our results, it could be hypothesized that the upregulation of *CCRL2*, which is more prominent among men with MDS, may be implicated in the gender-related differences in chronic myeloid diseases. The evaluation of a possible role of AR in the regulation of *CCRL2* in malignant myeloid cells is required to address this question.

CCRL2 is upregulated in a subset of malignancies including prostate (*34*) and colon (*35*) cancer. In contrast, downregulation of CCRL2 promotes lung cancer progression in animal models, apparently through suppressed immune surveillance (*36*). To our knowledge, the role of CCRL2 in myeloid malignancies has not been previously studied.

CCRL2 is critical for the activation of IL-8/CXCR2 signaling in neutrophils (*18*), and this pathway has been implicated in the growth of MDS cells via AKT and ERK signaling (*19*). Our analysis highlighted a more prominent role of CCRL2 in JAK2/STAT regulation. This pathway induces AML cell proliferation (*37*), while STAT3 inhibition impairs the growth of MDS/AML cells *in vitro* and *in vivo* (*38*). MDS92 and MDS-L cells are IL-3 dependent (*21*). This cytokine signals through CD123 promoting JAK2 auto-phosphorylation (*39*), leading to receptor phosphorylation attracting STAT proteins that then get phosphorylated and transduce the signal to the nucleus(*39, 40*). The exact mechanisms affecting JAK2 auto-phosphorylation and induction of STAT phosphorylation are not fully understood (*39*), but various molecules including protein tyrosine phosphatases (*41*), and LNK adaptor protein (*42*) are known regulators of these events. Similarly, the JAK2V617F mutation suppresses the inhibitory effect of the JAK2 pseudokinase domain (JH2) leading to JAK2 dimerization and constitutive activation of JAK2/STAT signaling(*40*). Given that CCRL2 promotes JAK2 auto-phosphorylation and STAT phosphorylation without affecting the interaction of JAK2 and CD123, it is possible that it intensifies the activity of the tyrosine kinase domain (JH1) of JAK2, which includes the tyrosine1007/1008 and catalyzes the STAT phosphorylation (*39, 40*). The decreased effect of CCRL2 in cells carrying the JAK2V617F mutation could be explained by the fact that these cells exhibit increased activity of JH1 at baseline. However, it is possible that other cell-specific alterations could be associated with this observation. Similarly, further study is needed for the understanding of the exact mechanism implicated in the interaction of CCRL2 with JAK2.

JAK2 inhibition has been routinely used in patients with JAK2 mutated myeloid malignancies (*4*). JAK2 inhibition by fedratinib shows promising efficacy in MDS cell lines and JAK2 wild-type erythroleukemia cells. CCRL2 knockdown increases the anti-proliferative and anti-clonogenic effect of fedratinib independent of the presence of a JAK2V617F mutation. JAK2 inhibition suppressed the de-phosphorylation of JAK2, leading to hyper-phosphorylation of its cryptic site involving tyrosine1007/1008(*29, 31*). This phenomenon has been associated with the JAK2 inhibitor withdrawal syndrome observed in a subset of patients with myelofibrosis (*29, 30*). Thus, it is possible that CCRL2 knockdown increases the efficacy of fedratinib by suppressing JAK2 auto-phosphorylation caused by JAK2 inhibition. Alternatively, the additive effect of CCRL2 knockdown to fedratitib efficacy could suggest that CCRL2 may be implicated in the regulation of other growth inducing pathways in MDS and AML.

In conclusion, our data suggest that CCRL2, a target gene of AR in normal granulopoiesis, is upregulated in stem and progenitor cells from MDS patients and influences the growth of MDS and AML cells. CCRL2 appears to promote JAK2/STAT signaling potentially by increasing the JAK2 auto-phosphorylation and its interaction with STAT proteins. Finally, CCRL2 suppression increased the efficacy of fedratinib in MDS and AML cells, independent of the presence of a JAK2V617F mutation.

## MATERIALS AND METHODS

### Patients and samples processing

Bone marrow samples were procured from bone marrow aspirations of patients with MDS, MDS/MPN and CD34+ AML, and normal marrow was obtained as excess material from the harvests of normal donors for allogeneic bone marrow transplantation. Specimens were collected by the Johns Hopkins Kimmel Cancer Center Specimen Accessioning Core. Appropriate informed consent was obtained from all donors before specimen collection in accordance with the Declaration of Helsinki and under a research protocol approved by the Johns Hopkins Institutional Review Board. CD34+ cell subsets were isolated using the CD34 MicroBead kit (Miltenyi Biotec) as previously described (*43*).

### Cell lines and reagents

Human MDS92 and MDS-L cells(*21, 22*) were a kind gift from Dr Daniel T. Starczynowski, University of Cincinnati(*24*). MDS92 and MDS-L cells were cultured in RPMI 1640 (Thermo Fisher Scientific, Waltham, MA) with 10% fetal bovine serum (FBS; MilliporeSigma, Burlington, MA), and 10 ng/ml IL-3 (PeproTech). OCI-AML3, DAMI, TF-1 Kasumi-1 and KG-1 cells were are purchased from ATCC and cultured in previously described conditions(*44–46*). All the cell lines were cultured with 2 mM L-glutamine, and 100 U/mL penicillin, and 100 μg/mL streptomycin at 37°C in 5% CO2. Primary CD34+ cells were cultured as previously described(*47*). Chlophosome-A chlodronate liposomes were purchased from FormuMax Scientific (#F70101C-A). Fedratinib was purchased from Selleckchem (TG101348) and was diluted in dimethyl sulfoxide (DMSO).

### CCRL2 knockdown

Lentiviral vectors expressing CCRL2-targeting short hairpin RNA (RHS3979-201740504, RHS39379-201740506) (Horizon), or empty pLKO.1-puro lentiviral vector were transfected together with pCMV-dR8.9 and vesicular stomatitis virus G–expressing plasmids into 293T cells using Lipofectamine 2000 (Thermo Fisher Scientific) for lentiviral supernatant production as previously described (*48*). MDS cells were incubated with the viral supernatant and 8 µg/mL polybrene (MilliporeSigma) for transduction. After at least 48 hours, cells were treated with 2 µg/mL of puromycin (MilliporeSigma) for 4 days to select for resistant cells.

### Flow cytometry analysis

Cells from healthy controls and MDS/AML patients were labeled with the following fluorescently-conjugated antibodies: 7AAD (BD Biosciences) (#559925), and Fluorescein isothiocyanate (FITC)-conjugated anti-CD34 (BD Biosciences) (#555821). Cells from AML patients were also labeled with BV510-conjugated anti-CD45 (BD Biosciences) (#5632204). CD34+CD38-cells from healthy controls and MDS patients cells were also labeled with the BV605-conjugated anti-CD38 (BD Biosciences) (#562666). Cells were then fixed in 4% formaldehyde for 15 minutes and permeabilized in 100% methanol for 30 minutes. Subsequently, cells were labeled with phycoerythrin (PE)-conjugated anti-CCRL2 (BioLegend) (#358303). For the analysis of STAT3/5 phosphorylation cells were also labeled with phycoerythrin (PE)-cyanine 7 (Cy7)-conjugated anti-phospho-STAT3 (Tyr705) (Thermo Fisher Scientific) (#25-9033-42) and allophycocyanin (APC)-conjugated anti-phospho-STAT5 (Tyr694) (Thermo Fisher Scientific) (#17-9010-42). For the assessment of CD11b expression MDS92 and MDS-L cells were labeled with the phycoerythrin (PE)-cyanine 7 (Cy7)-conjugated CD11b (BioLegend) (#557743). For the assessment of CD71 expression in TF-1 and DAMI cells were labeled with the phycoerythrin (PE)-cyanine 7 (Cy7)-conjugated CD71 (BioLegend) (#113811). Gating was based on clearly distinguishable populations, or in the absence of such, the negative antibody control as previously described (*43*). For the assessment of apoptosis following treatment with azacitidine, cells were stained for Annexin V and PI (BD Biosciences) (#556547) as previously described (*49*). After staining, cells were analyzed using the BD LSR II (BD Biosciences). Median fluorescence intensity (MFI) was determined for each marker using FlowJo analysis software version 10.0.8 (FlowJo, Ashland, CO, USA).

### Q-RT-PCR

Quantitative real-time polymerase chain reaction (PCR) was performed as previously described(*48*). In brief, total RNA was extracted by using the RNeasy Mini Kit (Qiagen, Valencia, CA), and complementary DNA was synthesized by using the ProtoScript kit (New England BioLabs). Quantitative real-time PCR was conducted by using sequence-specific primers (CCRL2 forward 5’-CTTCGAGAAAAACGTCTC-3’ and reverse 5’-CATCATATTCATCCTCTGGTG-3’, DNMT1 forward 5’-CGTAAAGAAGAATTATCCGAGG-3’ and reverse 5’-GTTTTCTAGACGTCCATTCAC-3’, PIM1 forward 5’-CTCTTCAGAATGTCAGGATC-3’ and reverse 5’-GGATGGTTCTGGATTTCTTC-3’, BCL2 forward 5’-GATTGTGGCCTTCTTTGAG-3’ and reverse 5’-GTTCCACAAAGGCATCC-3’, MCL1 forward 5’-TAGTTAAACAAGAGGCTGG-3’ and reverse 5’-ATAAACTGGTTTTGGTGGTG-3’, MYC forward 5’-TGAGGAGGAACAACAAGATG-3’ and reverse 5’-ATCCAGACTCTGACCTTTTG-3’ and GAPDH forward 5’-CTTTTGCGTCGCCAG-3’ and reverse 5’-TTGATGGCAACAATATCCA-3’) (Sigma-Aldrich), the Radian SYBR Green Lo-ROX qPCR Kit (Alkali Scientific, Fort Lauderdale, FL), and the CFX96 real-time PCR detection system (Bio-Rad). The RNA expression was normalized based on GAPDH expression.

### Western blotting

For western blotting analysis protein extraction was performed as previously described (*50*). For western blotting analysis of the effects of CCRL2 knockdown and azacitidine treatment, cells were treated with 0.5 μΜ of azacitidine after 4 days of selection in 2 μg/ml of puromycin. Antibodies against P-JAK2 (Tyr1007/1008) (#3776), P-STAT3 (Tyr705) (#9131), STAT3 (#4904), P-AKT (Ser473) (#9271), AKT (#4691), P-ERK1/2 (Thr202/Tyr204) (#9101), ERK1/2 (#4695), DNMT1 (#5032), c-MYC (#5605), PIM1 (#3247), BCL2 (#4223), MCL-1 (#5453) and β-actin (#4970) were purchased from Cell Signaling Technology. Antibodies against JAK2 (sc-294), P-STAT5 (Tyr694) (sc11761) and STAT5 (sc-835) were purchased from Santa Cruz and antibody against CCRL2 (LS C382506) was purchased LSBio.

### Co-Immunoprecipitation

MDS-L cells were deprived of IL-3 for 48 hours. Cells were lysed before and after 30 minutes of 20 ng/ml IL-3 treatment as described above(*50*). Cell lysates were incubated overnight with sepharose bead conjugate JAK2 monoclonal antibody (Cell Signaling) (#4089) or sepharose bead conjugate isotype control (Cell Signaling) (#3423). Beads were then washed extensively and boiled with 30 μ

### Colony formation assay

Clonogenic assays were performed as previously described (*48*). Briefly, cells following treatment were collected, counted, and resuspended at a density of 2000 cells/mL in methylcellulose-based media prepared as previously described (*48*). After 10 to 14 days of incubation at 37°C in 5% CO2, the recovery of colony-forming units was determined by colony counting under bright-field microscopy. A cell aggregate composed of >50 cells was defined as a colony.

### MTT assay

MDS92 and MDS-L cells were plated in 96 well plates (15000 cells/well). After 72 hours of treatment with various doses of fedratinib, the 3-(4,5-dimethylthiazol-2-yl)-2,5-diphenyl-2H-tetrazolium bromide (MTT) assay (11465007001, Roche Diagnostics, Mannheim, Germany) was performed according to manufacturer’s instructions. Absorbance was measured at 570 nm (reference 750 nm). Cell viability was calculated by 100 x (Absorbance of Sample/Average absorbance of untreated control) for the respective dose for each cell line.

### MDS xenograft studies

MDS-L cells were transduced with shControl and shCCRL2 lentiviral shRNA and selected under 2 μg/ml puromycin. Resistant cells were transduced with a retroviral vector carrying an enhanced green fuorescent protein (GFP) firefly luciferase fusion gene(*51*). Subsequently GFP+ cells were selected by cell sorting and injected intravenously to 10-week-old NOD.Cg-Prkdc^scid^ Il2rg^tm1Wjl^ Tg(CMV-IL3,CSF2,KITLG)1Eav/MloySzJ male mice (The Jackson Laboratory, stock no. 013062) (10^6^ cells/mouse) 48 hours after intraperitoneal injection of Clophosome-A clodronate liposomes (100 μl/mouse) as previously described(*25*). Bioluminescence signal was measured by using IVIS spectrum in vivo imaging system at days 1, 12, 22, 34, 55, 69 and 77. Peripheral blood was collected from the mice at days 24, 55 and 78 after the injection. At day 78, the mice were sacrificed and the percentage of human CD45+ cells was assessed in peripheral blood, spleen and bone marrow by flow cytometry.

### Publically available database

The analysis of mRNA levels of AR-target genes in normal stem and progenitor cells and various types of AML and MDS was based on RNA-sequencing data from the BloodSpot database (211434_s_at) (*20*).

### Statistical analysis

Statistical analyses were performed by using GraphPad Prism (GraphPad Software, La Jolla, CA). R programming language was used to visualize the mRNA expression levels of AR-target genes in healthy stem and progenitor cells and MDS and AML cells. Linear regression was performed to compare the mRNA and protein levels of CCRL2, and to assess the correlation of the MFI of CCRL2 with the MFI of P-STAT3/STAT5 in CD34+ cells from MDS and AML patients. Otherwise, an unpaired 2-tailed Student t test was performed to evaluate statistical significance.

## COMPETING INTERSTS

CLB has patents/patent applications in the fields of T-cell and/or gene therapy for cancer. CLB has received research funding from Merck, Sharpe, and Dohme, Kiadis Pharma, and Bristol Myers Squibb. The rest of the authors have no disclosures or conflicts of interest to report.

## AUTHOR CONTRIBUTIONS

TK and RJJ conceived and designed the study and wrote the manuscript. TK and BP performed the flow cytometry analysis. TK, EH and BWD performed the lentiviral transduction experiments. TK, EH and TR performed the western blot analysis. TK, PT, IC, CB and GG performed the xenograft studies. TK, BP, BCP and CE processed the primary samples. TK and CE performed the clonogenic assays. TK and PT performed the Q-RT-PCR analysis. TK and ARM performed the analysis of publicly available databases. TK and RV performed the statistical analysis. BWD, CB, ARM, LPG and MJL interpreted the data and edited the manuscript.

## SUPPLEMENTARY FIGURES

**Supplementary Figure 1. A.** Gating strategy for CD34+ and CD34+CD38-cells from healthy controls and MDS patients based on SSC-A/FSC-A and 7AAD. **B.** Gating strategy for CD34+ blasts from patients with AML based on SSC-A/FSC-A, and SSC-A/CD45. **C.** CCRL2 expression in representative samples of CD34+ cells from MDS and MDS/MPN patients in comparison to healthy controls **D.** No differences were observed in the CCRL2 protein levels expressed by CD34+ cells from patients with different subtypes of MDS (MDS with <5% blasts, MDS with >5% blasts and MDS/MPN). **E.** CD34+CD38-cells from men with MDS express higher levels of CCRL2 at the protein levels compared to CD34+CD38-cells from women with MDS (P=0.008), n=3 women and n=7 men. Graph shows the mean value and standard deviation.

**Supplementary Figure 2. A.** Western blotting showing the protein levels of CCRL2 in de novo AML (OCI-AML3, Kasumi-1), sAML (KG-1, DAMI) and MDS (MDS92 and MDS-L) cell lines. **B**. CCRL2 was knocked down in MDS92 and MDS-L by transduction with 2 different shRNAs (sh1 and sh2) targeting CCRL2 using empty vector as control. The efficacy of the knockdown was assessed at the RNA and protein level. The RNA expression of CCRL2 was suppressed in MDS92 cells (*P*<0.001 with both sh1 and sh2) and MDS-L cells (*P*=0.004 with sh1 and *P*=0.002 with sh2). The protein expression of CCRL2 was suppressed in MDS92 cells (*P*=0.001 with sh1 and *P*=0.008 with sh2) and MDS-L cells (*P*=0.001 with sh1 and *P*=0.002 with sh2), n=3. **C.** The protein levels of CCRL2 is significantly lower in MDS-L cells transduced with shCCRL2 compared to MDS-L cells transduced with shControl (*P*=0.006) following selection to puromycin for 10 days. **D.** Hemoglobin (Hgb), platelets (PLT), total white blood cells (WBC) and absolute neutrophil counts (ANC) in mice injected with MDS-L cells transduced with shControl and shCCRL2 at 24, 55 and 78 days after the injection. **E.** The percentage of human CD45+ cells is higher in the spleens of mice injected with MDS-L cells transduced with shControl and shCCRL2 (*P*=0.012), n=5. Graphs show the mean value and standard deviation.

**Supplementary Figure 3. A.** Representative western blotting showing the effect of CCRL2 knockdown by two different shRNAs (sh1 and sh2) in the phosphorylation of AKT (Ser473) and ERK1/2 (Thr202/Tyr204) in MDS92 and MDS-L with empty vector (shControl) as control. **B.** No significant differences were noted in the phosphorylation of AKT (Ser473) and ERK1/2 (Thr202/Tyr204). CCRL2 knockdown significantly decreases the phosphorylation of JAK2 (Tyr1007/1008), STAT3 (Tyr105) and STAT5 (Tyr694) in MDS92 and MDS-L cells, n=3. **C.** Representative western blotting showing the effect of CCRL2 knockdown on the protein levels of the JAK2/STAT target genes: MYC, PIM1, BCL2, MCL1 and DNMT1. **D.** The relative protein levels of the JAK2/STAT target genes: MYC, BCL2, PIM1, MCL1 and DNMT1 in MDS92 and MDS-L transduced with shControl and shCCRL2 shRNAs. CCRL2 knockdown suppresses the protein levels of the JAK2/STAT target genes in both MDS cell lines, n=3. Graphs show the mean value and standard deviation.

**Supplementary Figure 4. A.** No significant differences were observed in the phosphorylation of STAT3 and STAT5 in CD34+ cells from healthy controls, MDS and AML patients, n=8 healthy controls, n=7 MDS, n=4 AML. **B.** The expression of CCRL2 was not correlated with the phosphorylation of STAT5 in CD34+ cells from MDS and AML patients (Coef 0.009, *P*=0.750). n=11. Graphs show the mean value and standard deviation.

**Supplementary Figure 5. A.** MTT assay for the assessment of the effect of fedratinib in various MDS and AML cell lines, n=3. **B.** The IC50 of fedratinib in various MDS and AML cell lines. MDS-L and MDS92 are the most sensitive cell lines to fedratinib, n=3.

## SUPPLEMENTARY TABLES

**Supplementary Table 1.** Characteristics of healthy individuals, MDS and MDS/MPN and AML patients.

|  | **Healthy individuals**<br>**(N=12)** | **MDS and**<br>**MDS/MPN (N=22)** | **AML (N=12)** |
| --- | --- | --- | --- |
| Median age (range) | 44.5 years (18 – 54 years) | 66.0 years (46 – 79 years) | 59.0 years (20 – 70 years) |
| Gender | Women (7)<br>Men (5) | Women (10)<br>Men (12) | Women (5)<br>Men (8) |
| Disease subtype |  | MDS <5% blasts (10)<br>MDS >5% blasts (7)<br>MDS/MPN (5) | De novo (7)<br>sAML (6) |

**Supplementary Table 2.** Comparison of the entire MDS and MDS/MPN cohort to the MDS patients included in the CD34+CD38-analysis.

|  | **Entire MDS and**<br>**MDS/MPN cohort** | **MDS patients included in**<br>**the CD34+CD38- analysis** | **P-value** |
| --- | --- | --- | --- |
| Median age (range) | 66.0 years (46.0 – 79.0) | 62.0 years (46.0 – 78.0 years) | 0.363 |
| Male gender | 12 (0.55) | 6 (0.6) | 0.781 |
| Median blasts<br>percentage (range) | 0.02 (0.01 – 0.20) | 0.04 (0.01 – 0.20) | 0.605 |
| Median IPSS-R (range) | 4 (1 – 7.5) | 5.25 (1 – 7.5) | 0.489 |

