## Supplementary figures and images for "The role of the atypical chemokine receptor CCRL2 in myelodysplastic syndrome and secondary acute myeloid leukemia"

### Supplementary Figure 1

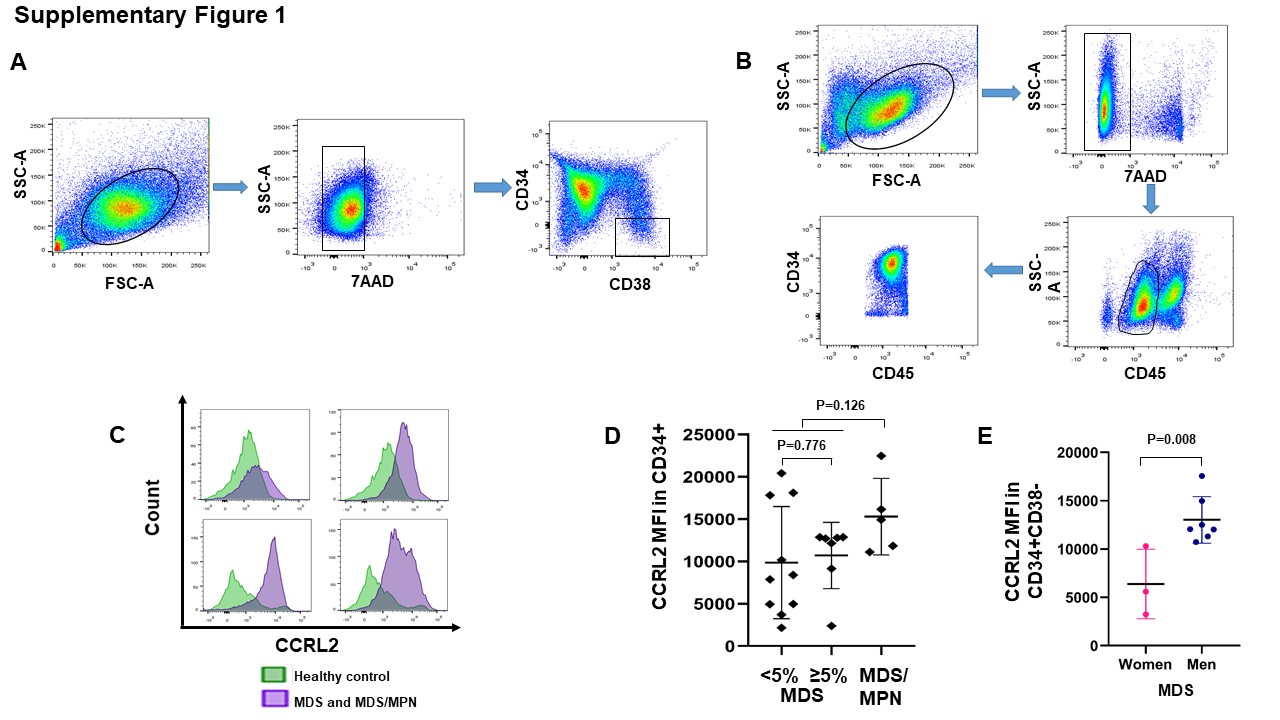

### Supplementary Figure 2

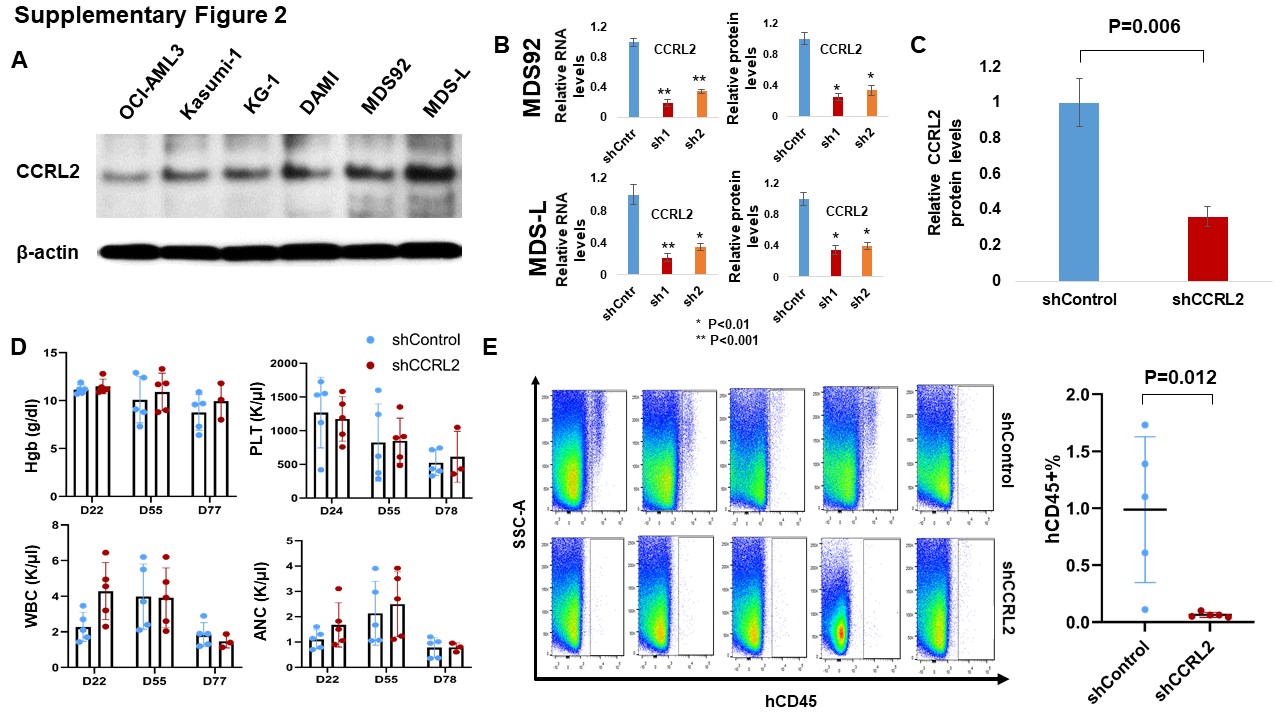

### Supplementary Figure 3

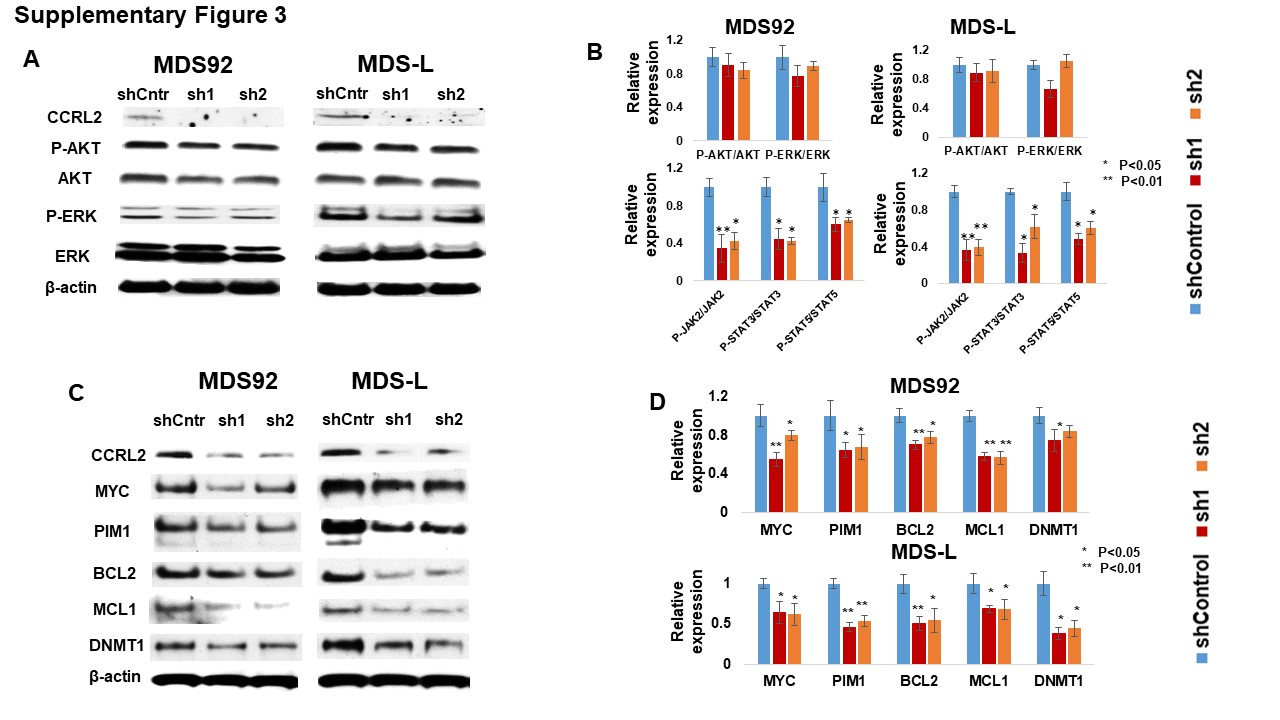

### Supplementary Figure 4

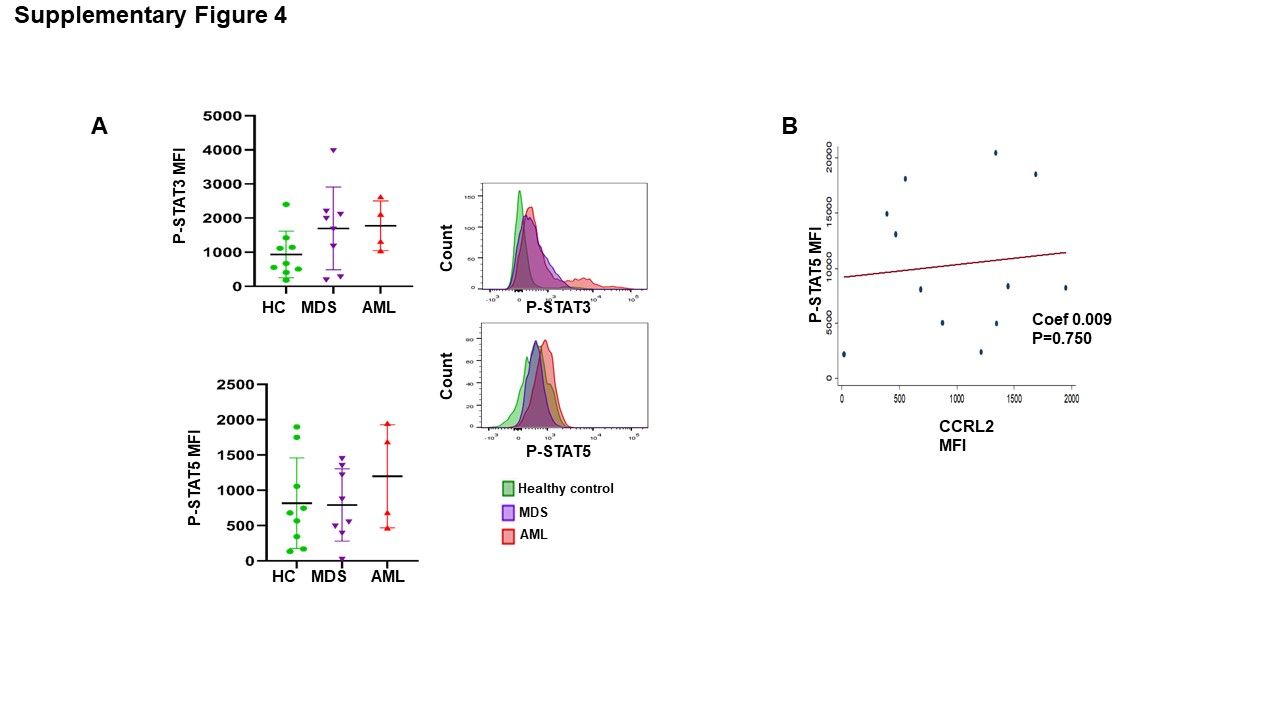

### Supplementary Figure 5

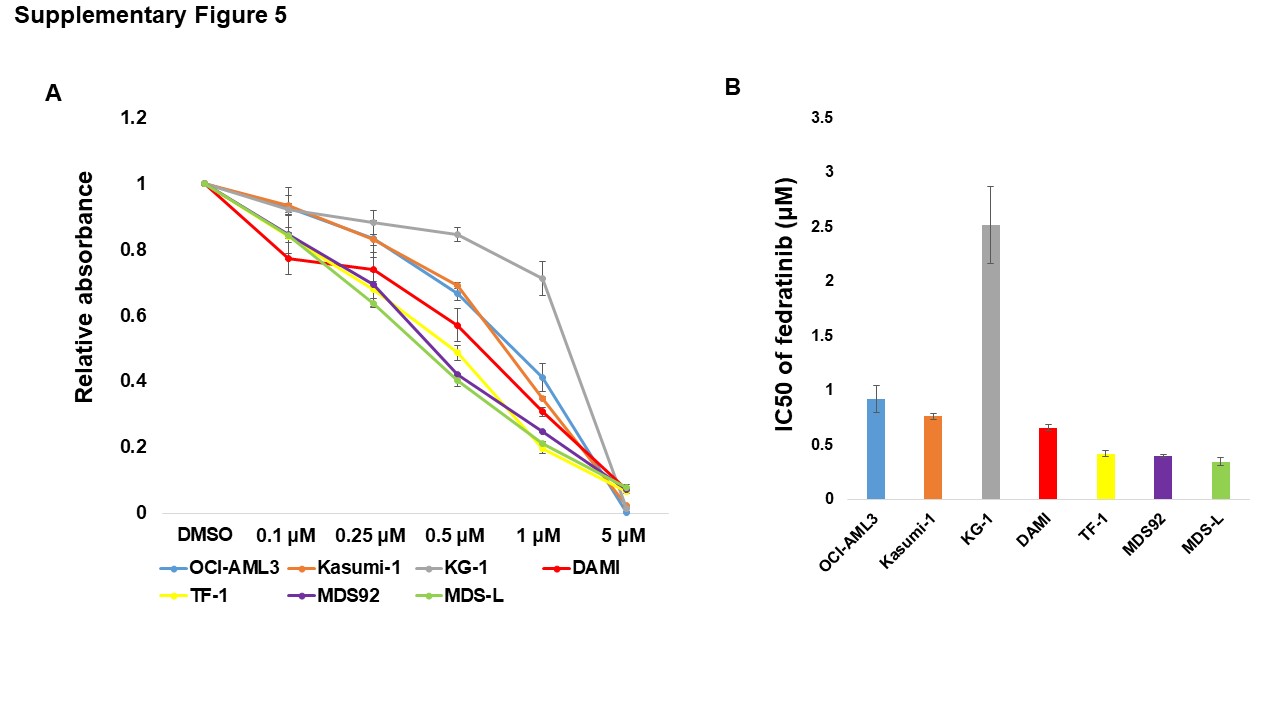
